# Age but not disease progression defines CD4^+^ and CD8^+^ T stem cell memory levels in human retroviral infections: contrasting effects of HTLV-1 and HIV-1

**DOI:** 10.1101/473363

**Authors:** Soraya Maria Menezes, Fabio Eudes Leal, Susan Pereira Ribeiro, Tim Dierckx, Mario Roederer, Devi SenGupta, Douglas F. Nixon, Johan Van Weyenbergh

## Abstract

**Background:** Human CD4^+^ and CD8^+^ stem cell memory T cells (T_SCM_) represent a minor fraction of circulating lymphocytes characterized by stemness and long-term *in vivo* persistence. CD4^+^ T_SCM_ are preferentially infected and constitute a reservoir for HIV-1, whereas CD8^+^ T_SCM_ appear to play a protective role. However, little is known about CD4^+^ and CD8^+^ T_SCM_ in the only other human pathogenic retroviral infection, human T-cell leukemia virus type 1 (HTLV-1). HTLV-1 is the etiological agent of both Adult T-cell Leukemia (ATL) and HTLV-1 associated myelopathy/tropical spastic paraperesis (HAM/TSP), a neuroinflammatory disorder. In ATL, CD4^+^ T_SCM_ cells were identified as the hierarchical leukemic stem cell, but data in HAM/TSP are lacking. Age is a major risk factor for both ATL and HAM/TSP, as both diseases generally manifest several decades after infection. Therefore, we explored a possible link between T_SCM_, age and disease status in human retroviral infections in a cross-sectional study, using multiparametric flow cytometry.

**Results:** We found that CD4^+^ or CD8^+^ T_SCM_ levels (quantified as CD3^+^CD45RA^+^CD45RO^−^ CD27^+^CCR7^+^Fas^hi^) do not differ between healthy controls and untreated HTLV-1 infected individuals with and without neuroinflammatory disorder. However, we found both T_SCM_ as well as CD8^+^ T_SCM_ significantly accumulated with age, resulting in a >400% increase in elderly HTLV-1 infected individuals (>60 years). A significant correlation between age and T_SCM_ signature genes was validated at the transcriptome level in an independent cohort. CD8^+^ but not CD4^+^ T_SCM_ were significantly decreased in untreated HIV-1 infection. Unexpectedly, CD8^+^ T_SCM_ recovery upon successful antiretroviral treatment was essentially complete (92.2±11.0%) in younger (<45 years) individuals, but significantly lower (37.3±6.1%) in older (>45 years) individuals (p=0.0003).

**Conclusion:** In HTLV-1 infection, an age-dependent accumulation of CD4^+^ and CD8^+^ T_SCM_ points towards a possible protective role of CD8 T_SCM_ in the elderly against leukemic but not neuroinflammatory disease. HIV-1-infected individuals lose their ability to restore CD8^+^ T_SCM_ levels upon successful antiretroviral therapy at later age (>45 years), which might eventually lead to immunological failure and decreased vaccine efficacy.

## Introduction

Human CD4^+^ and CD8^+^ stem cell memory T cells (T_SCM_) represent a small fraction (2-3%) of circulating lymphocytes characterized by intrinsic apoptosis resistance, proliferation and long-term *in vivo* persistence (Gattinoni et al., 2011). CD4^+^ T_SCM_ are preferentially infected and constitute a long-lived cellular reservoir for human immunodeficiency virus-1 (HIV-1) (Buzon et al., 2014), whereas CD8^+^ T_SCM_ appear to play a protective role (Ribeiro et al., 2014; Vigano et al., 2015). However, little is known about CD4^+^ and CD8^+^ T_SCM_ in the only other human pathogenic retroviral infection, namely human T-cell leukemia virus type 1(HTLV-1). HTLV-1 is the etiological agent of both Adult T-cell Leukemia (ATL) and HTLV-1 associated myelopathy/tropical spastic paraperesis (HAM/TSP), a neuroinflammatory disorder. Age is a major risk factor for both ATL and HAM/TSP, as both diseases generally manifest several decades after infection. In ATL, CD4^+^ T_SCM_ cells were identified as the hierarchical leukemic stem cell (Nagai et al., 2015). In HAM/TSP, we recently documented a predominant Fas^hi^ phenotype linked to lymphoproliferation and inflammation (Menezes et al., 2017), but its possible link to CD4^+^ and CD8^+^ T_SCM_ subsets is unknown. Interestingly, *FAS* polymorphisms determine both ATL susceptibility (Farre et al., 2008) and CD4^+^ and CD8^+^ Tscm levels in a large twin study (Roederer et al., 2015). Pulko et al. recently identified a novel human memory T cell subset with a naïve phenotype, accumulating with age (Pulko et al., 2016). While Pulko et al. elaborately explored various CD8^+^ naïve and memory cellular subsets and elegantly described a CD45RA^+^IFN-γ^+^CXCR3^+^Fas^Io^ subset of memory T cells with a naïve phenotype (T_MNP_) and responsiveness to chronic but not acute viral infections, their study did not include T memory stem cells (T_SCM_) (Pulko et al., 2016). Therefore, we explored a possible link between T_SCM_, age and disease status in human retroviral infections in a cross-sectional study.

## Results and Discussion

We found that CD4^+^ or CD8^+^ T_SCM_ levels (quantified as CD3^+^CD45RA^+^CD45RO^−^ CD27^+^CCR7^+^Fas^hi^) do not differ between healthy controls (HC, seronegative for HIV-1 and HTLV-1) and untreated HTLV-1 infected individuals with and without neuroinflammatory disorder (AC and HAM/TSP patients, Fig. 1A). Whereas the hallmark of HIV-1 infection is CD4^+^ depletion, HTLV-1 infection propels CD4^+^ cells into the cell cycle while protecting CD8^+^ cells from apoptosis (Sibon et al., 2006). However, CD4^+^ T_SCM_ abundance was not significantly different between HC, untreated HTLV-1 (HIV-1 negative), untreated HIV-1 and treated HIV-1 infection (successful antiretroviral therapy with undetectable viral load), as shown in Fig. 1B. Nevertheless, CD8^+^ T_SCM_ was significantly decreased in untreated HIV-1 infection compared to all other groups (Fig. 1C). In parallel to HTLV-1 infection, CD4^+^ and CD8^+^ T_SCM_ levels (%) did not differ in untreated HIV-1-infected individuals with different degrees of disease progression, i.e. controllers, non-controllers and immunological progressors (Ribeiro et al., 2014).

**Figure 1.**
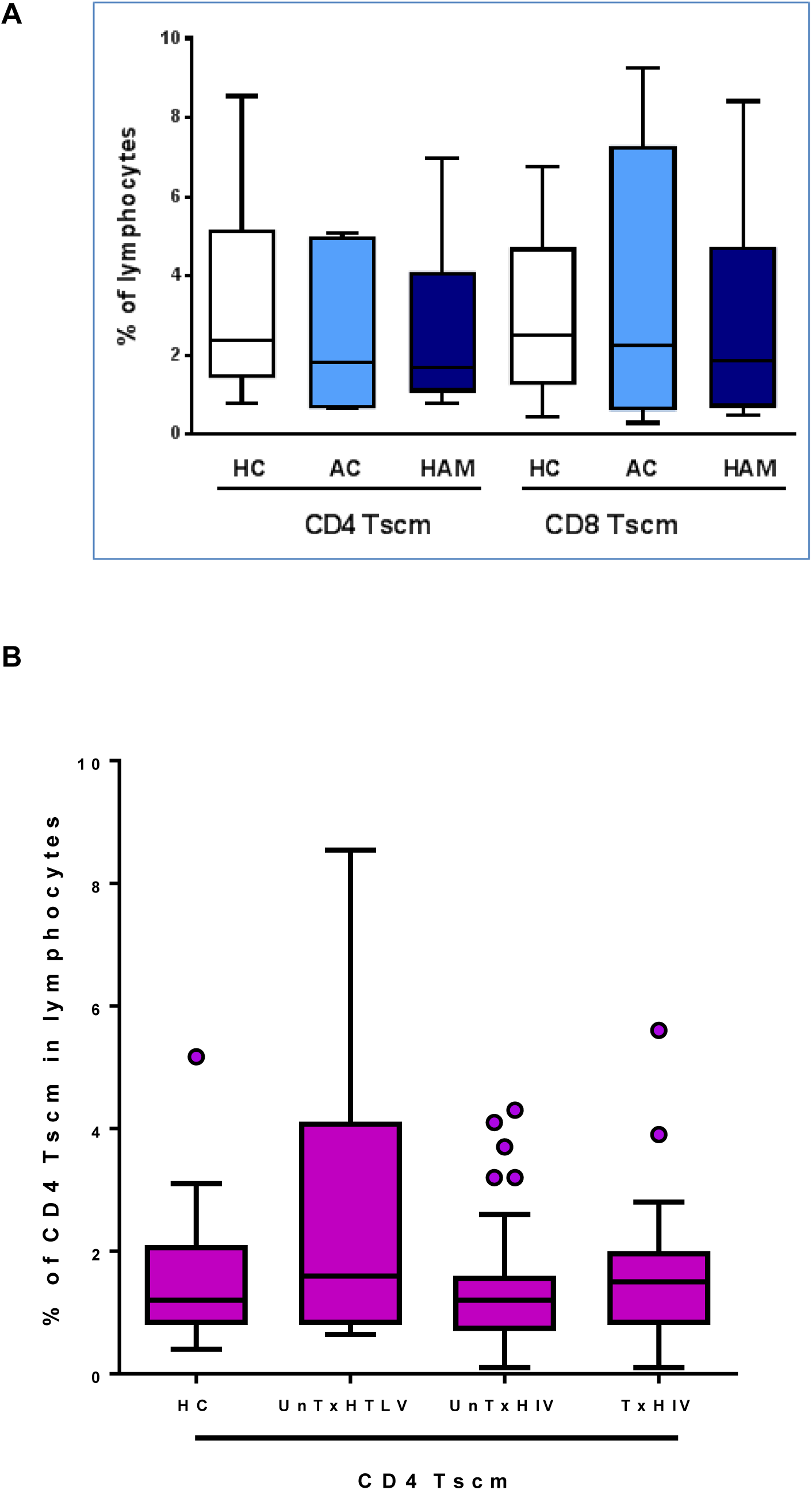
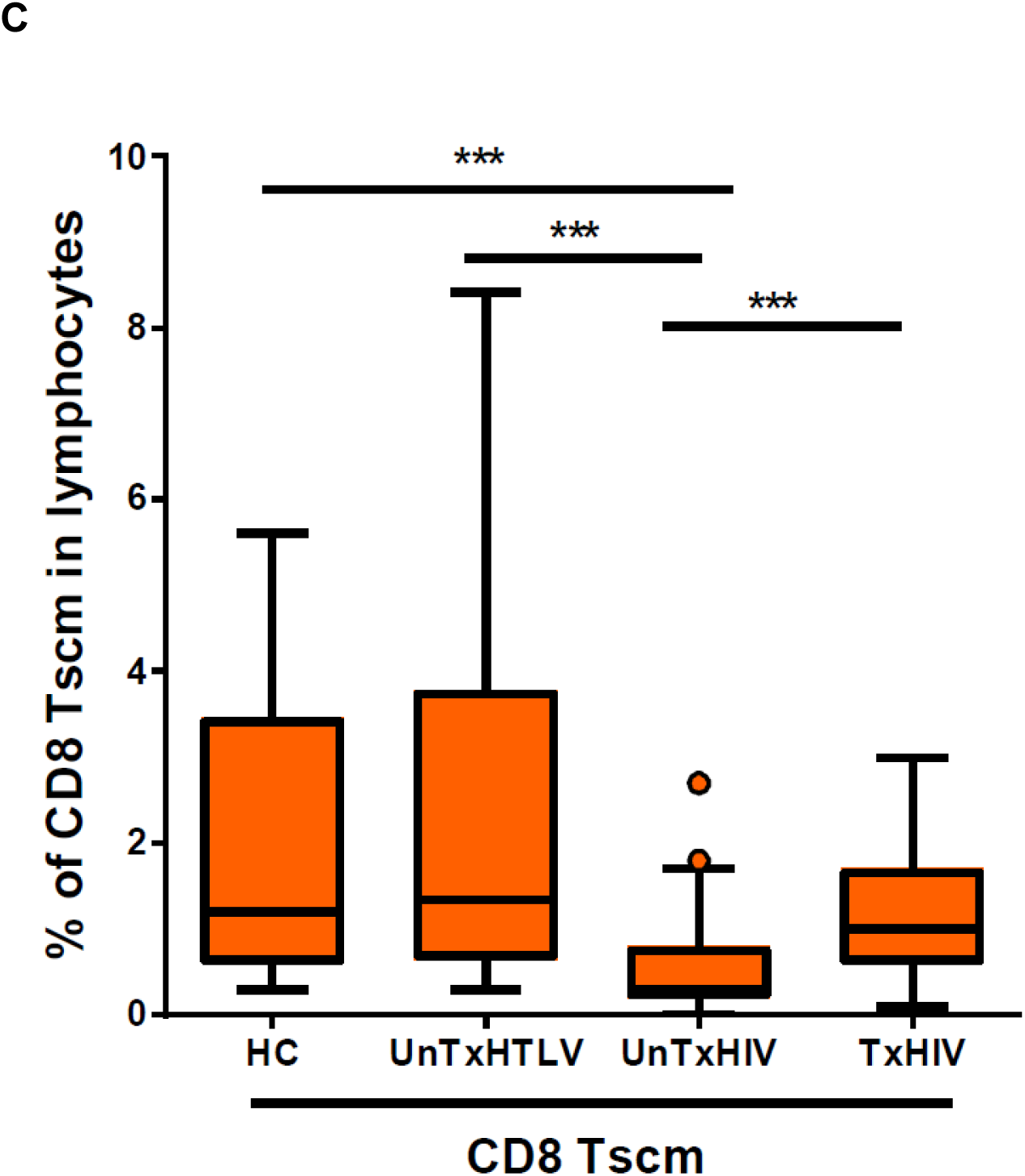
CD4^+^ and CD8^+^ T_SCM_ levels do not differ between matched healthy controls (HC), HTLV-1 asymptomatic carriers (AC) and HAM/TSP patients, while CD8^+^ T_SCM_ cells are depleted in untreated HIV-1-infected individuals. (A) No significant difference in the levels of CD4^+^ and CD8^+^ T_SCM_ cells (CD3^+^CD45RA^+^CD45RO^−^CD27^+^CCR7^+^Fas^hi^) quantified by flow cytometry in HC (n=7 CD4^+^ T_SCM_, n=9 CD8^+^ T_SCM_), AC (n=6) and HAM/TSP patients (n=9). (B) No significant difference in the levels of CD4 ^+^ T_SCM_ between HC (n=19), untreated HTLV-1- (n=13), untreated HIV-1- (n=77) and treated (n=27) HIV-1-infected individuals. (C) CD8^+^ T_SCM_ levels are decreased in untreated HIV-1-infected individuals when compared to HC, untreated HTLV-1 and treated HIV-1 infected individuals. (***p < 0.001; Kruskal-Wallis test with Dunn’s post-test).

In a large cohort of healthy controls (n=460), CD4^+^ T_SCM_ were found to modestly increase by 25% over a 30-year period in HC (p=0.06, Roederer et al. unpublished), whereas CD8^+^ T_SCM_ levels decrease 25% over the same period (p=0.004, Roederer et al., unpublished). Unexpectedly, in untreated HTLV-1-infected individuals, both CD4^+^ T_SCM_ as well as CD8^+^ T_SCM_ significantly accumulated with age (r=0.60, p=0.016 and r=0.73, p=0.0019, respectively), resulting in a >400% increase in the elderly (>60 years Fig. 2A). This effect was not due to differences in viral factors, as CD4^+^ and CD8^+^ T_SCM_ levels did not correlate to proviral load nor viral mRNAs (Tax and HBZ). Furthermore, age-dependent accumulation was observed in the combined HTLV-1-infected group, as well as in the asymptomatic and HAM/TSP subgroups, implying that age, but not disease status, determines CD4^+^ T_SCM_ and CD8^+^ T_SCM_ levels in HTLV-1 infection.

**Figure 2:**
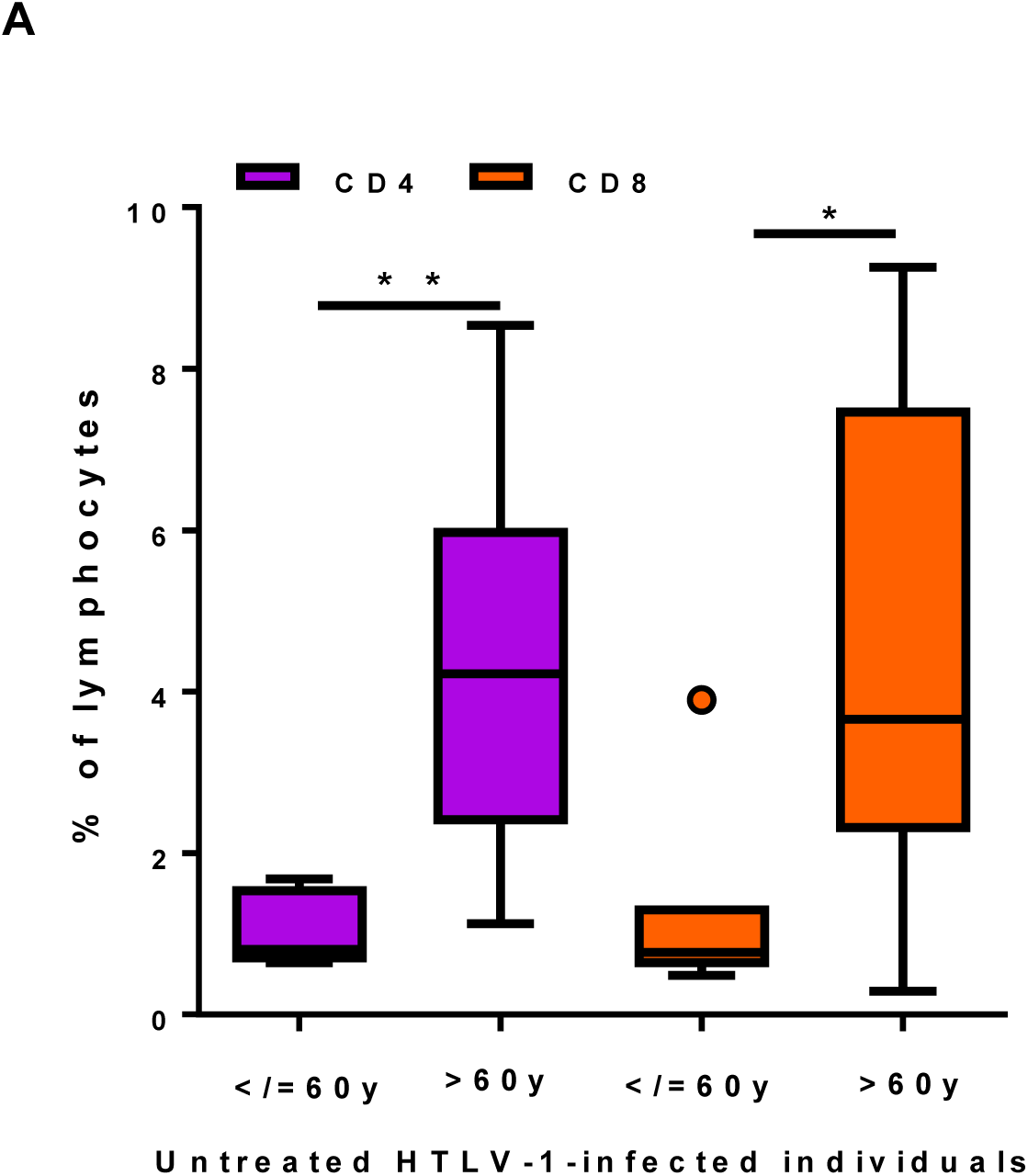
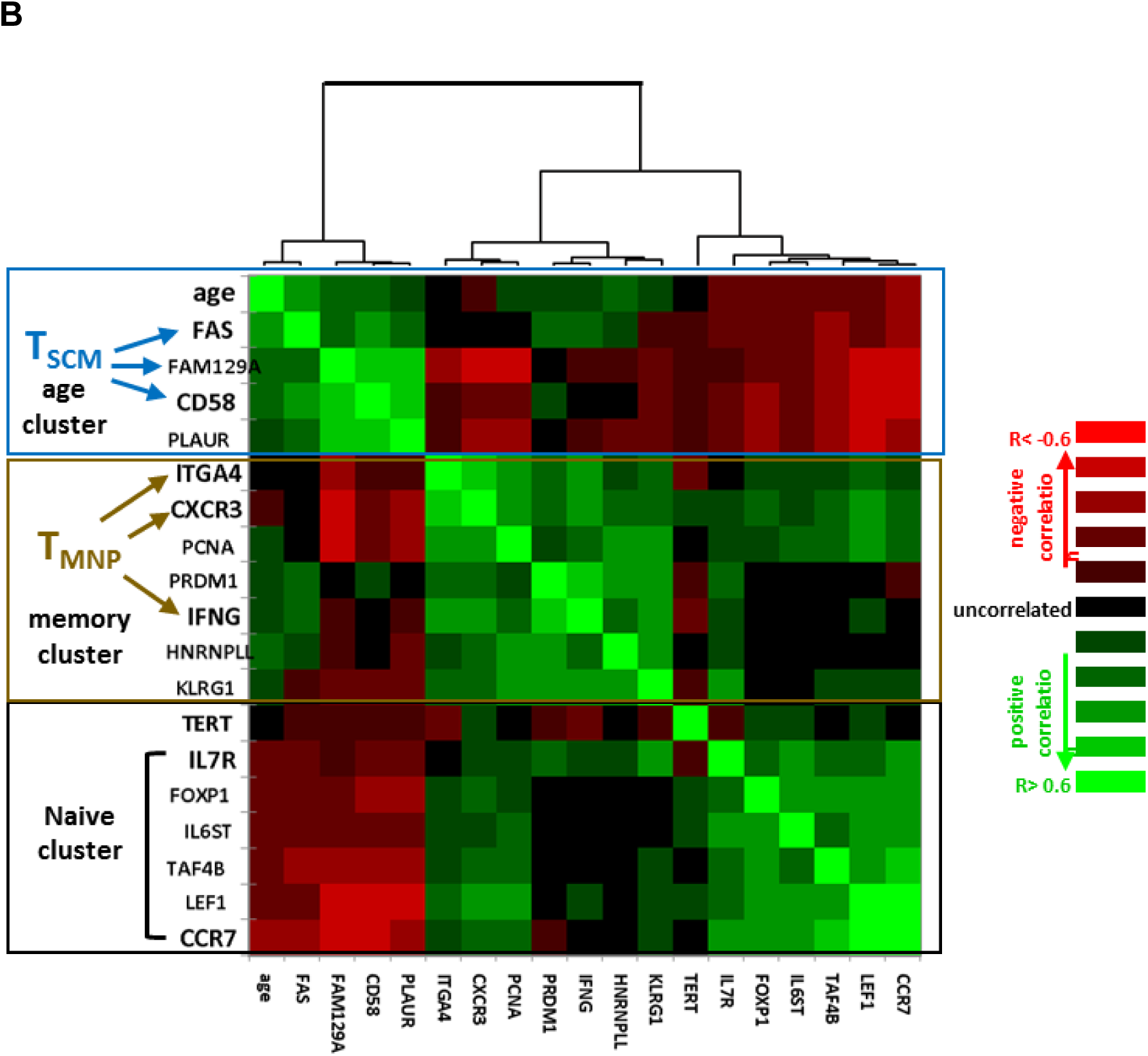
T_SCM_ cells accumulate with age and T_SCM_ but not T_MNP_ signature genes correlate to age in untreated HTLV-1 infection. (A) CD4^+^ (**p = 0.0024, unpaired t test) as well as CD8^+^ (*p = 0.016) T_SCM_ levels increase in older HTLV-1 infected individuals (> 60 years, n=9 vs. <60 years, n=7).(B) Transcriptomic analysis of T_SCM_, T_MNP_ and Naïve/Memory markers, clustered according to Spearman’s correlation (heatmap) reveals three distinct clusters: T_SCM_-Age, T_MNP_-Memory and Naïve, in HTLV-1 infected individuals from an independent published UK cohort (n=30).

We validated these findings at the transcriptome level using an independent, previously published cohort (Tattermusch et al., 2012). Transcript levels of T_SCM_ signature markers (Gattinoni et al., 2011) *FAS* (p=0.0005) and *CD58* (p=0.026) were positively correlated to age but not disease in HTLV-1-infected individuals (n=30), forming a distinct cluster with other T_SCM_ signature genes (*FAM129A* and *PLAUR*, upper cluster Fig. 2B). On the other hand, transcript levels of T_MNP_ signature genes *ITGA4*, *CXCR3* and *IFNG* were not correlated to age or disease status but significantly correlated to each other (p<0.001, in agreement with Pulko et al.), forming a second cluster (middle in Fig. 2B), which included memory (*PRDM1/HNRNPLL/KLRG1*) and proliferative (*PCNA*) markers. A third distinct cluster, negatively correlated to age, was comprised of “classical” naïve T cell marker genes (*IL7R, FOXP1, IL6ST, LEF1, TAF4B, CCR7*) and telomerase (*TERT*). Supporting HTLV-1 specificity, no naïve or memory subset marker genes were correlated to age in transcriptomes from HC (data not shown). This age-dependent increase is quite surprising given the role of CD4^+^ T_SCM_ in ATL pathogenesis (Nagai et al., 2015). Since the oldest HTLV-1-infected individuals (both AC and HAM/TSP) did not develop ATL, even at a mean age of 66±4.5 years, the parallel increase of CD8^+^ T_SCM_ cells might keep the pre-leukemic CD4^+^ T_SCM_ in check, in agreement with a proposed protective role of CD8^+^ cytotoxic effector cells in ATL (Rowan et al., 2016), which can be derived from the CD8^+^ T_SCM_ pool. A wealth of recent studies has indeed demonstrated the superior anti-tumoral properties and clinical potential of CD8^+^ T_SCM_ cells for adaptive immunotherapy, as compared to other CD8^+^ naïve or memory subsets.

In contrast to HTLV-1-infected individuals, we observed a significant age-dependent decline of CD8^+^ T_SCM_ (p=0.0017, r=-0.57) but not CD4^+^ T_SCM_ levels in treated HIV-1 patients (Fig. 3A). There was no correlation of CD4^+^ T_SCM_ or CD8^+^ T_SCM_ with age in untreated HIV-1 patients (Fig. 3B), again independent of disease status (data not shown). When calculating the percentage of CD8^+^ T_SCM_ depletion (relative to HC), CD8^+^ T_SCM_ depletion was strikingly similar between younger (<45 years) and older (>45 years) untreated HIV-1-infected individuals (Figure 3C). However, the percentage of CD8^+^ T_SCM_ recovery upon successful treatment was essentially complete (92.2±11.0%) in younger (<45 years) individuals, but significantly lower (37.3±6.1%) in older (>45 years) individuals (p=0.0003, Figure 3D), which parallels HIV-associated CD8^+^ senescence in this age group (Cobos Jimenez et al., 2016). Thus, contrary to HTLV-1 infection, disruption of CD8^+^ T_SCM_ homeostasis occurs at an early age in HIV-1 infection, but is relatively stable over the age groups (defective CD8 T_SCM_ recovery is apparent at different cut-offs, i.e. in age groups >40, >50 and >60 years, data not shown).

**Figure 3:**
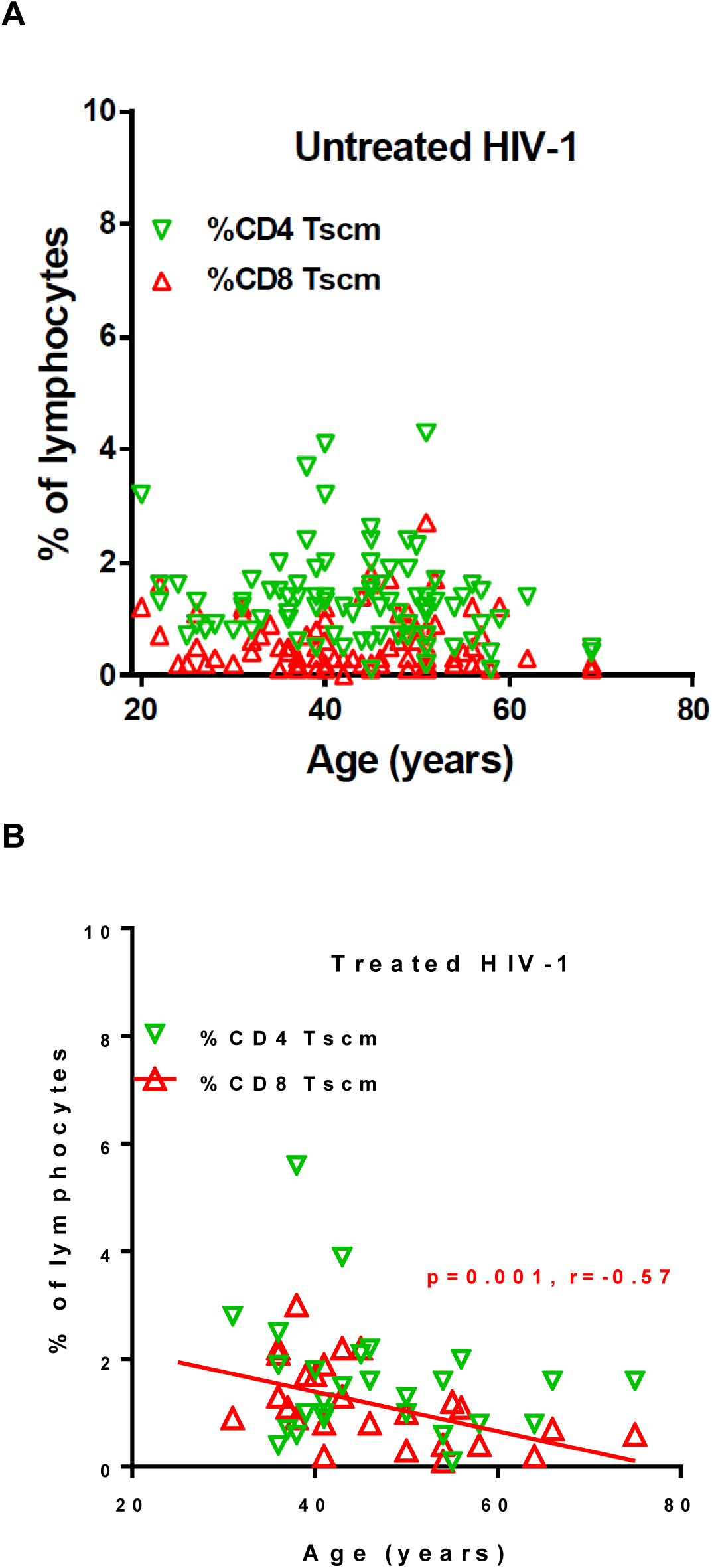
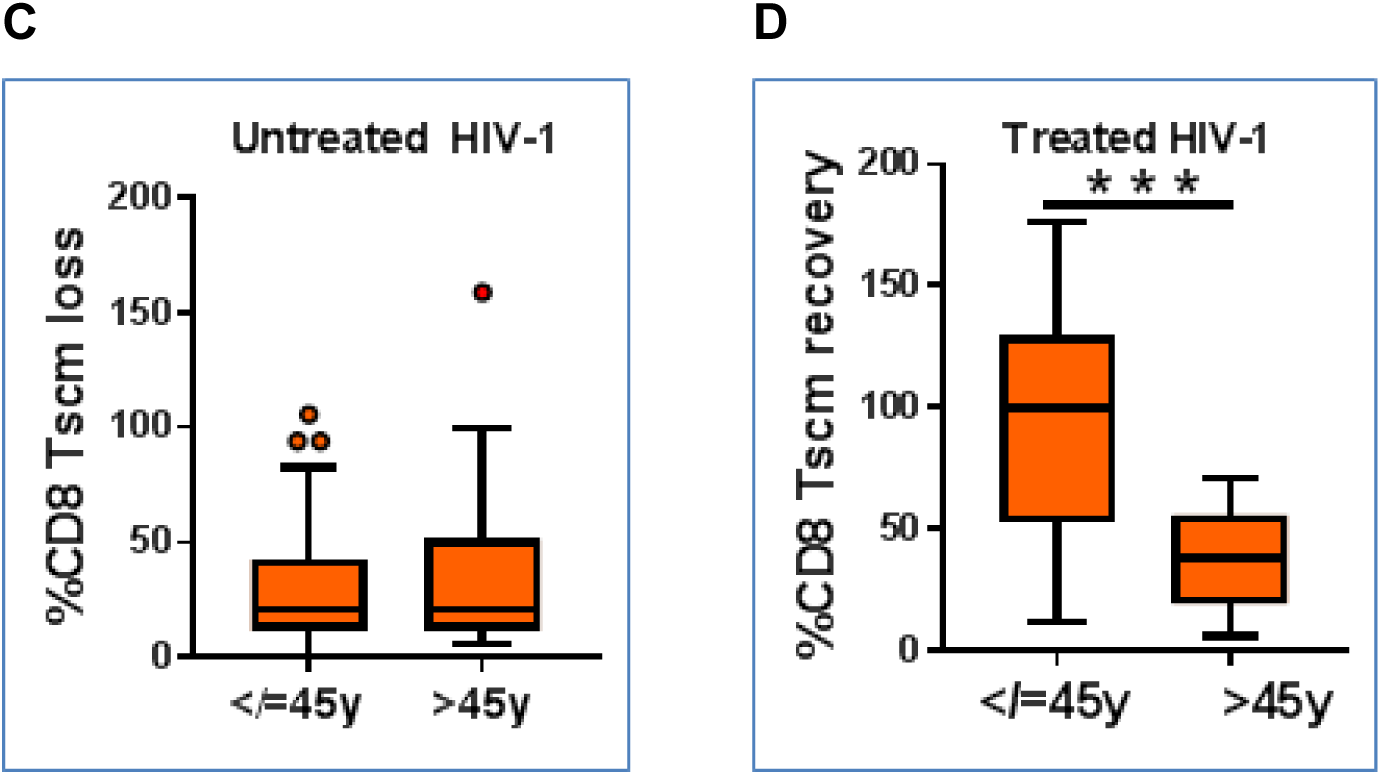
HIV-1 infected individuals over 45 years lose their ability to restore CD8^+^ T_SCM_ levels following successful ART. (A) In treated HIV-1-infected individuals, CD8^+^ but not CD4^+^ T_SCM_ levels negatively correlate with age. (**p=0.001, Spearman’s r=0.57, n=27). (B) Neither CD4^+^ nor CD8^+^ T_SCM_ levels were correlation of age to in untreated HIV-1-infected individuals (n=77). (C) No age effect in CD8^+^ T_SCM_ depletion (relative to matched HC) in untreated HIV-1 infected individuals (> and < 45 years). (D) HIV-1 infected individuals on successful ART treatment >45 years recover less than half of CD8^+^ T_SCM_ levels (relative to matched HC) compared to those <45 years. (***p<0.001, Unpaired t test, with Welch correction)

Hence, our findings have obvious clinical implications in the light of current life-long requirement of antiviral therapy for HIV-1-infected individuals. Although successful as measured by suppressed viral load and increased CD4^+^ levels, antiretroviral therapy was unable to restore protective CD8^+^ T_SCM_ levels at middle age. This eventually might lead to immunological failure and disease progression, highlighting the need for long-term in-depth immunological follow-up of large cohorts, beyond the current routine CD4^+^ and viral load monitoring. In addition, this novel age-T_SCM_ link should be considered in ongoing therapeutic vaccination trials aiming at a functional cure for HIV. On a larger scale, given the extensively documented role of CD8^+^ T_SCM_ in life-long immunological memory and an increasingly ageing human population, natural as well as vaccine-induced herd immunity against other known as well as “novel” pathogens (e.g. pandemic flu, arboviruses) might wane or disappear over time, considering the 25% decline observed in a large cohort of healthy controls. In conclusion, our findings indicate vigilance is warranted in the light of age-dependent disruption of T_SCM_ homeostasis.

## References

Buzon, M.J., Sun, H., Li, C., Shaw, A., Seiss, K., Ouyang, Z., Martin-Gayo, E., Leng, J., Henrich, T.J., Li, J.Z., et al. (2014). HIV-1 persistence in CD4+ T cells with stem cell-like properties. Nature medicine 20, 139–142.

Cobos Jimenez, V., Wit, F.W., Joerink, M., Maurer, I., Harskamp, A.M., Schouten, J., Prins, M., van Leeuwen, E.M., Booiman, T., Deeks, S.G., et al. (2016). T-Cell Activation Independently Associates With Immune Senescence in HIV-Infected Recipients of Long-term Antiretroviral Treatment. The Journal of infectious diseases 214, 216–225.

Farre, L., Bittencourt, A.L., Silva-Santos, G., Almeida, A., Silva, A.C., Decanine, D., Soares, G.M., Alcantara, L.C., Jr., Van Dooren, S., Galvao-Castro, B., et al. (2008). Fas 670 promoter polymorphism is associated to susceptibility, clinical presentation, and survival in adult T cell leukemia. J Leukoc Biol 83, 220–222.

Gattinoni, L., Lugli, E., Ji, Y., Pos, Z., Paulos, C.M., Quigley, M.F., Almeida, J.R., Gostick, E., Yu, Z., Carpenito, C., et al. (2011). A human memory T cell subset with stem cell-like properties. Nature medicine 17, 1290–1297.

Menezes, S.M., Leal, F.E., Dierckx, T., Khouri, R., Decanine, D., Silva-Santos, G., Schnitman, S.V., Kruschewsky, R., Lopez, G., Alvarez, C., et al. (2017). A Fas(hi) Lymphoproliferative Phenotype Reveals Non-Apoptotic Fas Signaling in HTLV-1-Associated Neuroinflammation. Frontiers in immunology 8, 97.

Nagai, Y., Kawahara, M., Hishizawa, M., Shimazu, Y., Sugino, N., Fujii, S., Kadowaki, N., and Takaori-Kondo, A. (2015). T memory stem cells are the hierarchical apex of adult T-cell leukemia. Blood 125, 3527–3535.

Pulko, V., Davies, J.S., Martinez, C., Lanteri, M.C., Busch, M.P., Diamond, M.S., Knox, K., Bush, E.C., Sims, P.A., Sinari, S., et al. (2016). Human memory T cells with a naive phenotype accumulate with aging and respond to persistent viruses. Nature immunology 17, 966–975.

Ribeiro, S.P., Milush, J.M., Cunha-Neto, E., Kallas, E.G., Kalil, J., Somsouk, M., Hunt, P.W., Deeks, S.G., Nixon, D.F., and SenGupta, D. (2014). The CD8(+) memory stem T cell (T(SCM)) subset is associated with improved prognosis in chronic HIV-1 infection. Journal of virology 88, 13836–13844.

Roederer, M., Quaye, L., Mangino, M., Beddall, M.H., Mahnke, Y., Chattopadhyay, P., Tosi, I., Napolitano, L., Terranova Barberio, M., Menni, C., et al. (2015). The genetic architecture of the human immune system: a bioresource for autoimmunity and disease pathogenesis. Cell 161, 387-403.

Rowan, A.G., Witkover, A., Melamed, A., Tanaka, Y., Cook, L.B., Fields, P., Taylor, G.P., and Bangham, C.R. (2016). T Cell Receptor Vbeta Staining Identifies the Malignant Clone in Adult T cell Leukemia and Reveals Killing of Leukemia Cells by Autologous CD8+ T cells. PLoS pathogens 12, e1006030.

Sibon, D., Gabet, A.S., Zandecki, M., Pinatel, C., Thete, J., Delfau-Larue, M.H., Rabaaoui, S., Gessain, A., Gout, O., Jacobson, S., et al. (2006). HTLV-1 propels untransformed CD4 lymphocytes into the cell cycle while protecting CD8 cells from death. The Journal of clinical investigation 116, 974–983.

Tattermusch, S., Skinner, J.A., Chaussabel, D., Banchereau, J., Berry, M.P., McNab, F.W., O'Garra, A., Taylor, G.P., and Bangham, C.R. (2012). Systems Biology Approaches Reveal a Specific Interferon-Inducible Signature in HTLV-1 Associated Myelopathy. PLoS pathogens 8, e1002480.

Vigano, S., Negron, J., Ouyang, Z., Rosenberg, E.S., Walker, B.D., Lichterfeld, M., and Yu, X.G. (2015). Prolonged Antiretroviral Therapy Preserves HIV-1-Specific CD8 T Cells with Stem Cell-Like Properties. Journal of virology 89, 7829–7840.

